## Supplementary Information for "TraDIS-validate: a method for curating ordered gene-replacement libraries"

**Supplementary Material**

**Primers that did not map to the BW25113 reference genome:**

ycgR primer:	TTTGAGCTTTTTCTGCTACCGTTTGCCCATCAATCAGCGTCGCTCCGGCG

|||||||||||||||||||||||||||||||||||||||||||||||

genome:	TTTGAGCTTTTTCTGCTACCGTTTGCCCATCAATCAGCGTCGCTCCGCGT

argI primer:	GATGG-TTTCC-G-CATCTT-ATAGCGATCAG--ATTATTTACTGAGCGTCGCGAC

|| ||||| | |||||| |||||||| || ||||||||||||||||||||||

genome:	TTGTATTTCCCGGCATCTTTATAGCGAT-AGCAATTATTTACTGAGCGTCGCGAC

lacI primer: TGAGCTAACTCACATTAATTGCGTTGCGCTCACTGCCCGCTTTCCAGTCG

||||||||||||||||||||||||||||||

genome: GCAGATGCGGTCCTCAATCGCGCGTTGCGCTCACTGCCCGCTTTCCAGTCG

=30 nt homology

eutA primer up: CCAAAAGAAGACGCGACCGCGACTAAAACCGAAGCGGAGGCACAATG

eutA primer dn:TGTGGTCTTTAGTTTCATAAGTCGTTCCCTCAGGAAGGAAATGCGAGTGA

eutA primer dn rev-c:	TCACTCGCATTTCCTTCCTGAGGGAACGACTTATGAAACTAAAGACCACA

|||||||||||||||||||||

eutA genome:	AATCACTCGCATTTCCTTCCTGAATGCGGTATTCGCCAGCGG

eutA primer dn rev-c: TCACTCGCATTTCCTTCCTGAGGGAACGACTTATGAAACTAAAGACCACA

|||||||||||||||||||||||||||||||||||||

eutB genome:	TCATCGATTCTTCCTGAGGGAACGACTTATGAAACTAAAGACCACA

The primer to construct the eutA Keio mutant has homology to the BW25113 reference genome in the absence of the CPZ-55 cryptic prophage.

**The artefact protein that arises when the *flu* mutant is constructed and cassette removed:**

>BW25113_Flu - artefact

M-MIPGIRRPAVRSSTSLGSIGTSKQLQPT-HTECDLLTEPSPLCGPGHHDRDPDRRNGSSTPHSLALTIT*

*Flu start codon – cassette scar – neighbouring ORF

**Construction of the mltC mutant using the primers reported in the Keio paper:**

>BW25113_2963 mltC membrane-bound lytic murein transglycosylase C 3097792:3098871 forward

ATG-AAA-AAA-TAT-CTC-GCG-CTG-GCT-TTG-ATT-GCG-CCG-TTG-CTC-ATC-TCC-TGT-TCG-ACG-ACC-AAA-AAA-GGC-GAT-ACC-TAT-AAC-GAA-GCC-TGG-GTC-AAA-GAT-ACC-AAC-GGT-TTT-GAT-ATT-CTG-ATG-GGG-CAA-TTT-GCC-CAC-AAT-ATT-GAG-AAC-ATC-TGG-GGC-TTC-AAA-GAG-GTG-GTG-ATC-GCT-GGT-CCT-AAG-GAC-TAC-GTG-AAA-TAC-ACC-GAT-CAA-TAT-CAG-ACC-CGC-AGC-CAC-ATC-AAC-TTC-GAT-GAC-GGT-ACG-ATT-ACT-ATC-GAA-ACC-ATC-GCC-GGG-ACA-GAA-CCT-GCC-GCG-CAT-TTG-CGC-CGG-GCA-ATT-ATC-AAA-ACG-TTA-TTG-ATG-GGT-GAC-GAT-CCG-AGT-TCG-GTC-GAT-CTC-TAT-TCC-GAC-GTT-GAT-GAT-ATT-ACG-ATT-TCG-AAA-GAA-CCT-TTC-CTT-TAC-GGT-CAG-GTG-GTG-GAC-AAC-ACC-GGG-CAG-CCG-ATT-CGC-TGG-GAA-GGT-CGC-GCA-AGC-AAC-TTC-GCG-GAT-TAT-CTG-CTG-AAA-AAC-CGT-CTG-AAG-AGC-CGC-AGC-AAC-GGG-CTG-CGT-ATC-ATC-TAC-AGC-GTC-ACC-ATT-AAC-ATG-GTG-CCG-AAC-CAC-CTT-GAT-AAA-CGT-GCG-CAC-AAA-TAT-CTC-GGC-ATG-GTC-CGC-CAG-GCG-TCA-CGG-AAA-TAT-GGC-GTT-GAT-GAG-TCG-CTG-ATT-CTG-GCA-ATT-ATG-CAG-ACC-GAA-TCT-TCC-TTT-AAC-CCG-TAT-GCG-GTC-AGC-CGT-TCC-GAT-GCG-CTG-GGA-TTA-ATG-CAG-GTG-GTA-CAA-CAT-ACT-GCC-GGG-AAA-GAT-GTG-TTC-CGC-TCG-CAG-GGG-AAA-TCC-GGC-ACG-CCG-AGC-CGC-AGT-TTC-TTG-TTT-GAT-CCT-GCC-AGC-AAT-ATT-GAT-ACC-GGC-ACC-GCG-TAT-CTG-GCG-ATG-CTG-AAC-AAT-GTT-TAT-CTC-GGC-GGA-ATT-GAT-AAC-CCA-ACA-TCG-CGG-CGT-TAT-GCC-GTC-ATC-ACC-GCC-TAT-AAC-GGC-GGC-GCA-GGC-AGC-GTG-CTG-CGA-GTC-TTT-TCG-AAT-GAT-AAG-ATT-CAG-GCT-GCC-AAT-ATT-ATT-AAC-ACC-ATG-ACG-CCG-GGC-GAT-GTT-TAT-CAG-ACG-CTG-ACG-ACC-CGC-CAT-CCC-TCT-GCG-GAA-TCT-CGC-CGT-TAT-CTT-TAT-AAA-GTG-AAT-ACC-GCG-CAA-AAA-TCC-TAC-CGC-CGC-CGA-TAA

NNN = region deleted by kanamycin resistance cassette

**Truncation of *hcaT* when the *csiE* mutant is constructed:**

>hcaT MFS (major facilitator superfamily) transporter - 1970265: 1971404 MW: 41593.21

MVLQSTRWLALGYFTYFFSYGIFLPFWSVWLKGIGLTPETIGLLLGAGLVARFLGSLLIAPRVSDPSRLISALRVLALLTLLFAVAFWAGAHVAWLMLVMIGFNLFFSPLVPLTDALANTWQKQFPLDYGKVRLWGSVAFVIGSALTGKLVTMFDYRVILALLTLGVASMLLGFLIRPTIQPQGASRQQESTGWSAWLALVRQNWRFLACVCLLQGAHAAYYGFSAIYWQAAGYSASAVGYLWSLGVVAEVIIFALSNKLFRRCSARDMLLISAICGVVRWGIMGATTALPWLIVVQILHCGTFTVCHLAAMRYIAARQGSEVIRLQAVYSAVAMGGSIAIMTVFAGFLYQYLGHGVFWVMALVALPAMFLRPKVVPSC

*12TM domains underlined

NNN = region deleted by kanamycin resistance cassette

NNN = TM domain (x12)

**Primers for amplification of the cassette-gDNA junction:**

K1F 5′-TCGCCTTCTTGACGAGTTCTTCTAATAAGG-3′

A1R 5′-GACTGGAGTTCAGACGTGTGCTCTTCCGATC-3′

K2F 5′-AATGATACGGCGACCACCGAGATCTACACTCTTTCCCTACACGACGCTCTTCCGATCTNNNNNNAAAGTATAGGAACTTCGAAGCAGCT-3′

*Underlined section has homology to the kanamycin resistance cassette

NNN = barcode

A2R – NEBNext Illumina Index primers were used
