## Supplementary Figures for "TraDIS-validate: a method for curating ordered gene-replacement libraries"

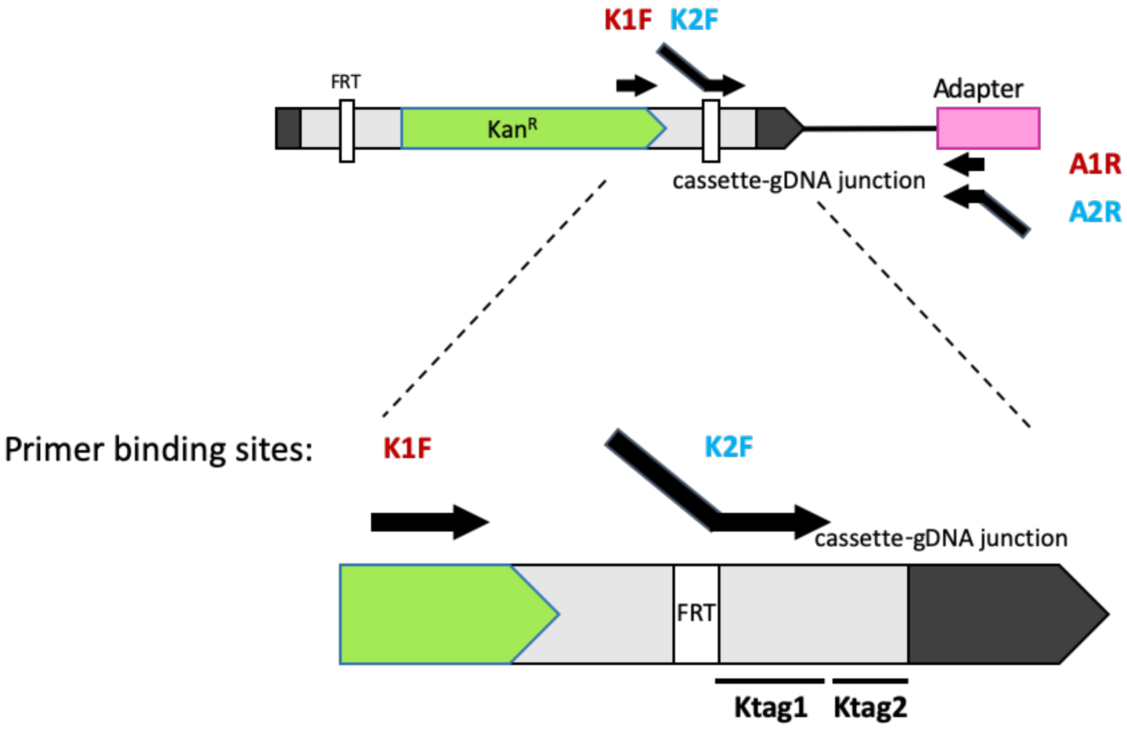


**Supplementary Figure 1. Primer binding sites for amplification of the cassette-gDNA junctions for sequencing**

Following DNA fragmentation, the cassette-gDNA junctions are enriched by PCR using primer pair K1F and A1R. The sample is then prepared for sequencing by PCR using the semi-nested primer pair K2F and the NEBNext® Multiplex Oligos for Illumina, denoted here as 'A2R'. These primers introduce the necessary barcodes and flow cell-binding adapters required for Illumina sequencing. During processing of FASTQ data, the kanamycin resistance cassette is recognized in two pattern matching steps that correspond with Ktag1, allowing for 3 mismatches, and Ktag2, allowing for 1 nucleotide mismatch. The Ktag1 sequence corresponds with the terminal 25 nucleotides of the K2F primer. FRT = Flp recognition target; gDNA = genomic DNA.

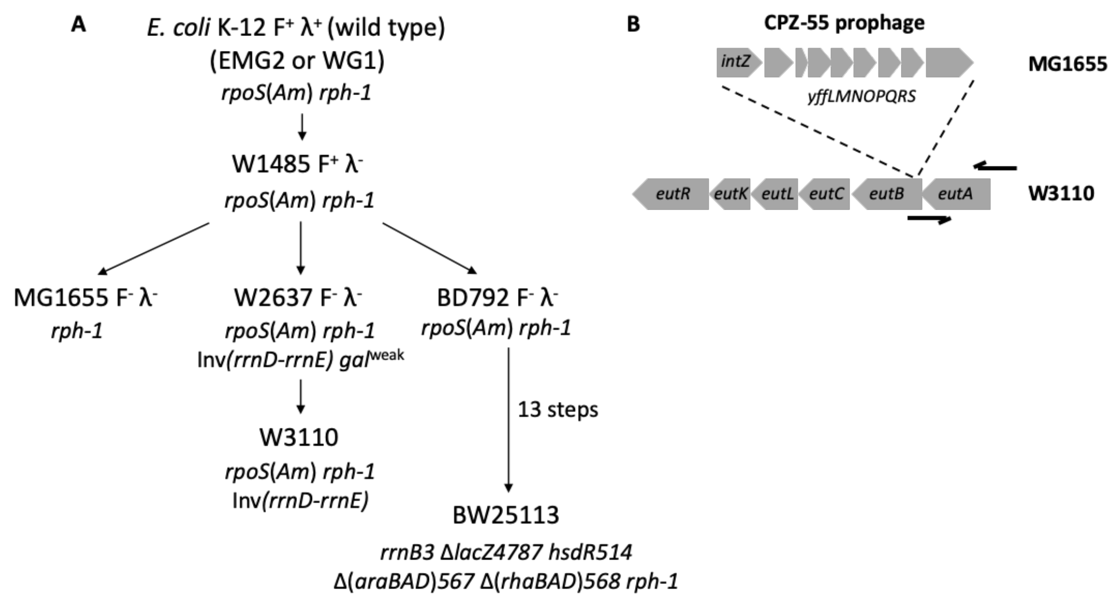


**Supplementary Figure 2. The relationship between *E. coli* K-12 strains BW25113, MG1655 and W3110**

(A) Figure adapted from Bachmann (1972), Baba *et al*. (2006) and Hayashi *et al.* (2006). (B) The *eut* operon of MG1655 and W3110 and the primer binding sites for the primers used to construct the *eutA* mutant.

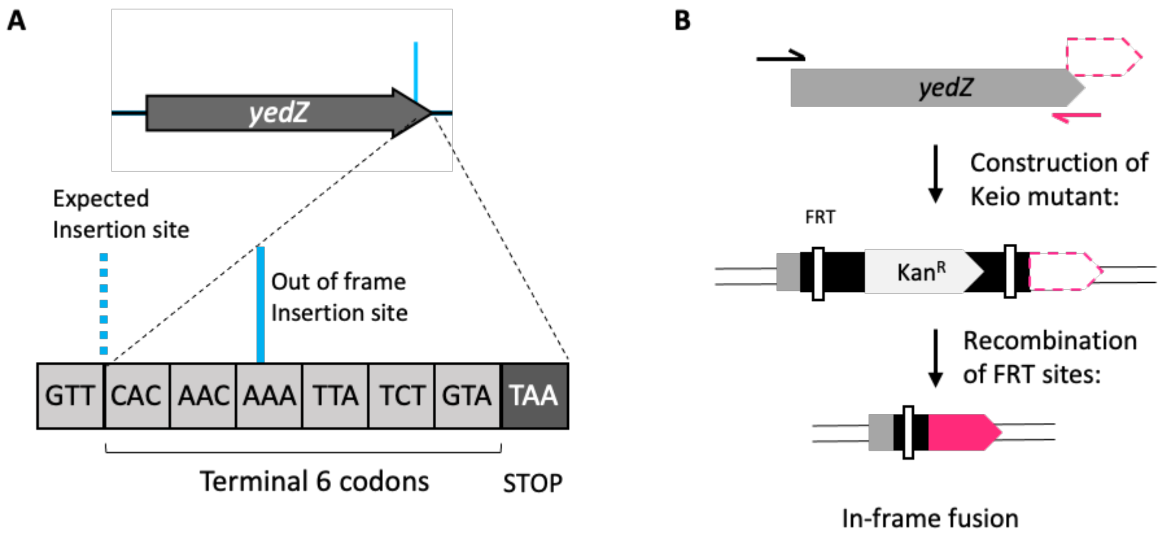


**Supplementary Figure 3. Identification of an out-of-frame insertion within *yedZ***

(A) Insertion of the kanamycin resistance cassette in the *yedZ* mutant. (B) Excising the kanamycin resistance cassette in the *yedZ* mutant will result in construction of an artefact coding sequence.

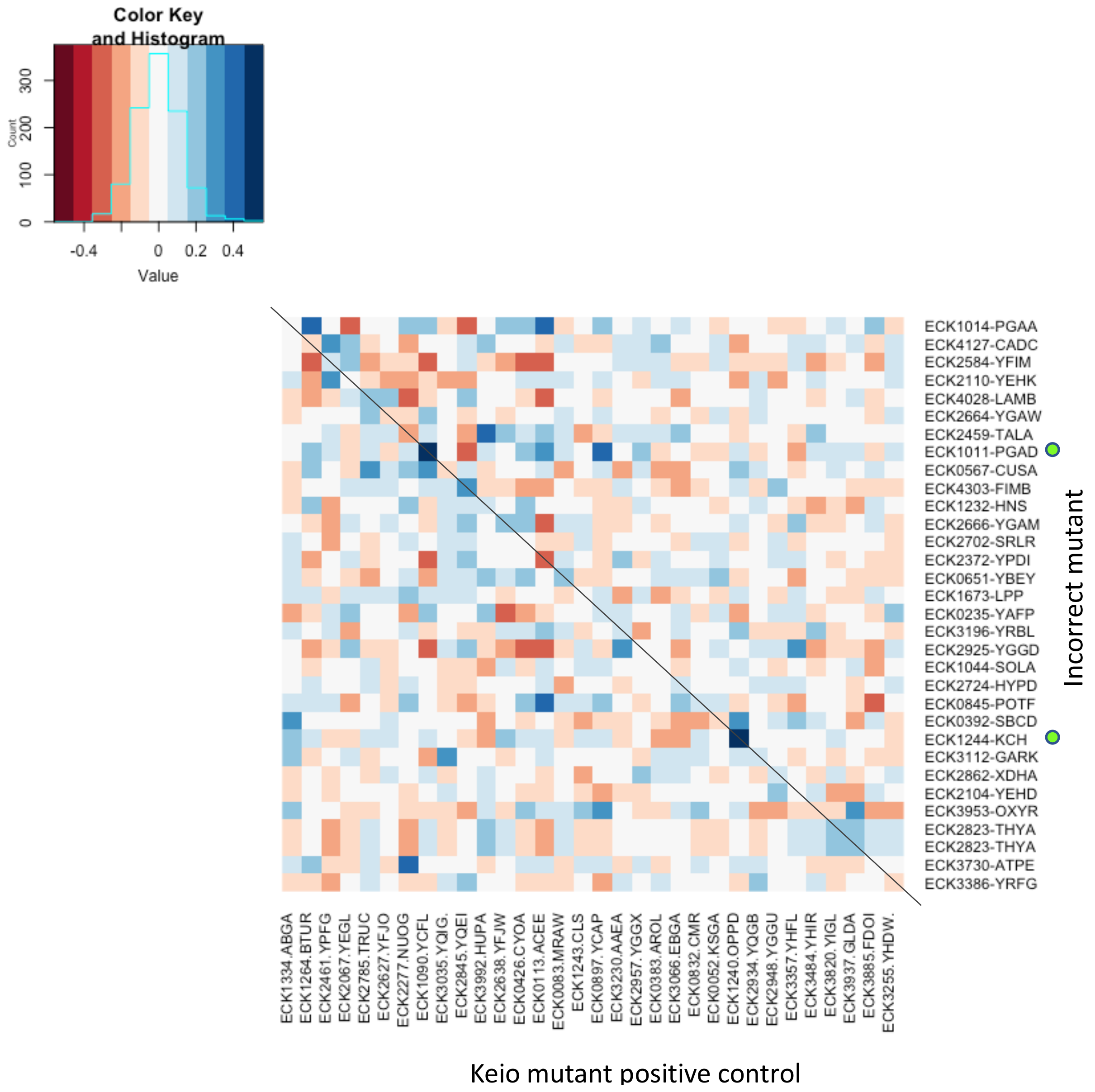


**Supplementary Figure 4. Comparison of the phenotypic profiles of Keio mutants with an incorrectly positioned kanamycin resistance cassette**

Comparison of phenotypic correlation scores, taken from Nichols *et al.* (2011). Mutants with an incorrectly located kanamycin resistance cassette (y-axis) were compared to the corresponding mutant (x-axis). Mutants with a negative phenotypic correlation are shown in red, mutants with a positive phenotypic correlation are shown in blue.
